# Prediction of protein subcellular localization using deep learning and data augmentation

**DOI:** 10.1101/2020.05.19.068122

**Authors:** Majid Ghorbani Eftekhar

## Abstract

Identifying subcellular localization of protein is significant for understanding its molecular function. It provides valuable insights that can be of tremendous help to protein’s function research and the detection of potential cell surface/secreted drug targets. The prediction of protein subcellular localization using bioinformatics methods is an inexpensive option to experimentally approaches. Many computational tools have been built during the past two decades, however, producing reliable prediction has always been the challenge. In this study, a Deep learning (DL) technique is proposed to enhance the precision of the analytical engine of one of these tools called PSORTb v3.0. Its conventional SVM machine learning model was replaced by the state-of-the-art DL method (BiLSTM) and a Data augmentation measure (SeqGAN). As a result, the combination of BiLSTM and SeqGAN outperformed SVM by improving its precision from 57.4% to 75%. This method was applied on a dataset containing 8230 protein sequences, which was experimentally derived by Brinkman Lab. The presented model provides promising outcomes for the future research. The source code of the model is available at https://github.com/mgetech/SubLoc.

## Introduction

The subcellular localization of proteins can supply essential information about their function and facilitate the study of complicated approaches that adjust biological procedures at the cellular level. That is why the prediction of protein subcellular localization (PSL) using bioinformatics methods is imperative. It plays a crucial role in the genome annotation task and verifying experimental results. The prediction of proteins on the cell surface of bacterial pathogens is also of high importance, because such proteins are likely to be primary drug or vaccine targets. Several attributes present within the protein’s primary structure can impact its subcellular localization, for example, the presence of signal peptide or membrane-spanning alpha-helices. Identification of PSL by experimental methods is a tedious and time-consuming task. That is why many different computational methods have been built to carry out this task during the past 20 years. Basically, prediction tools take, for instance, a protein sequence of amino acids as input and predict its location inside the cell as the outcome.

One of the programs that was originally developed for predicting protein localization in prokaryotic organisms (Gram-negative bacteria) is POSRT by Kenta Nakai [1]. PSORT was developed into a set of programs (PSORT, PSORT II, iPSORT) potential of predicting proteins from all classes of organisms. PSORTb v3.0 [2] is an updated version of PSORT algorithm that is being developed by the Brinkman Laboratory [3], which offers more advanced analytical models based on a dataset of experimentally derived localizations. Many other methods concentrate on enhancing prediction accuracy—maximizing the number of positive predictions on the training dataset, at the expense of producing more false positive results. On the other hand, PSORTb v3.0 is designed to emphasize precision (or specificity) over accuracy and recall (or sensitivity). It is leveraging varied methods as part of its analytical engine such as SCL-BLAST, Support Vector Machines (SVM) [4], ModHMM (an extension of PRODIV-HMM [5]). It is believed that SVM is a conventional Machine Learning (ML) method and to improve the PSORTb’s precision, adopting the more state-of-the-art methods such as Deep Learning (DL) techniques can be beneficial. In this study with an effort to increase the PSORTb’s overall precision, the performance of the SVM algorithm in prediction of PSL is compared against a DL technique called BiLSTM, which also uses SeqGAN [6] as a Data augmentation approach to compensate the lack of adequate protein sequences in certain classes (particularly the ones with multiple localization sites).

## Materials and methods

### Dataset

The dataset [7] of experimentally verified sequences containing 11,692 proteins was handpicked by the Brinkman Lab and made available to the public in FASTA format. The dataset of prokaryotic organisms contains Gram-negative, Gram-positive bacteria and Archaea that is consisted of 8230, 2652 and 2652 instances respectively. In this investigation, a subset of the dataset that contains protein sequences from Gram-negative bacteria was used. The classification of data according to their subcellular location include 5025 Cytoplasmic proteins, 50 Cytoplasmic/Cytoplasmic Membrane (Cyt/CM) proteins, 1607 Cytoplasmic Membrane proteins, 58 Cytoplasmic Membrane/Periplasmic (CM/Per) proteins, 437 Periplasmic proteins, 3 Periplasmic/Outer Membrane (Per/OM) proteins, 541 Outer Membrane proteins, 90 Outer Membrane/Extracellular (OM/Ext) proteins and 419 Extracellular proteins.

### Preprocessing

Protein sequences contain amino acid letters and in order for Machine Learning algorithms to be able to perform calculation on the them, they need to be encoded into numbers. An ordinal encoding approach, as shown in Figure 1, was adopted to convert each amino acid letter to ordinal integer codes.

**Figure 1.**
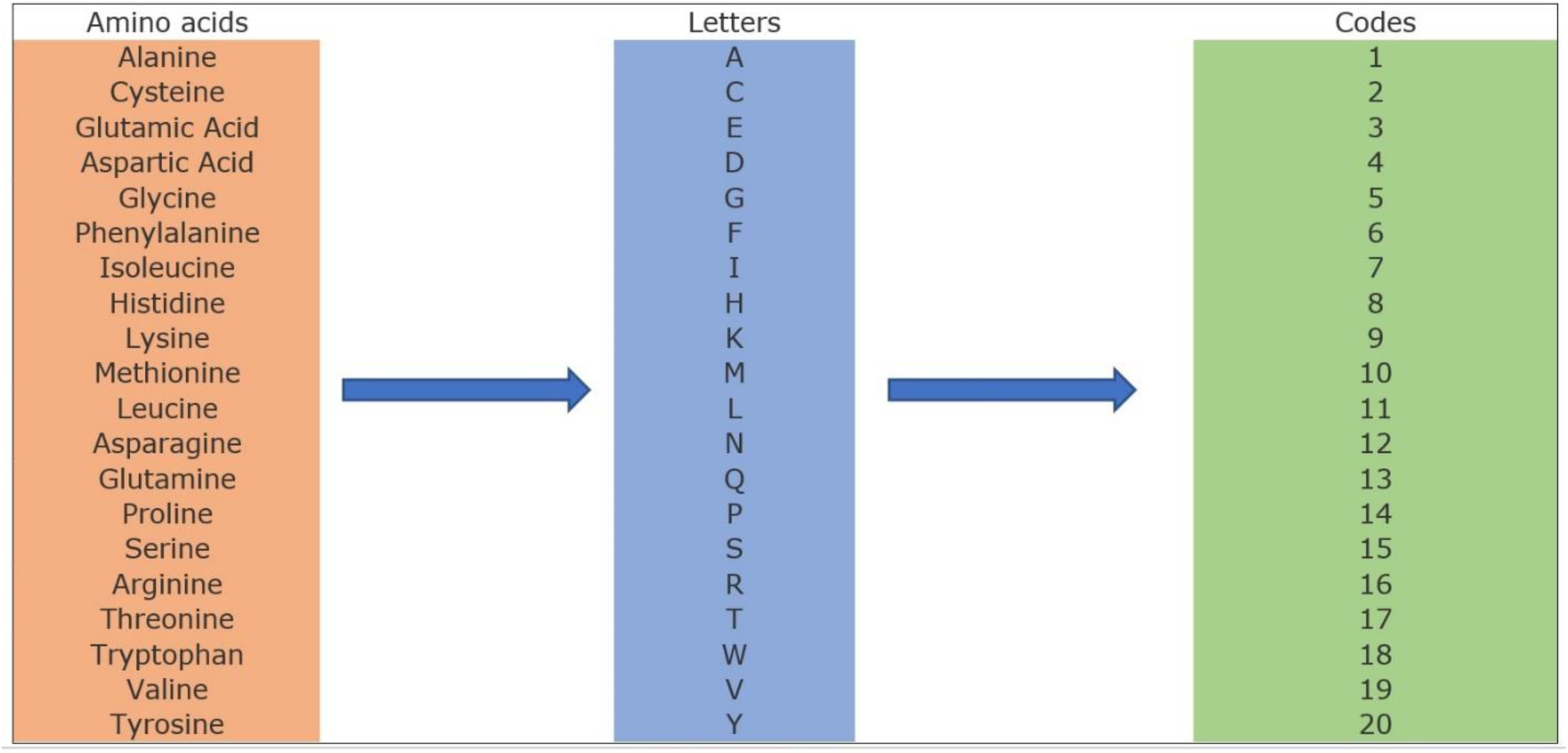
Amino acid encoder.

Feature selection is the process of reducing the number of input variables when developing a predictive model. It is desirable to reduce the number of input variables to lower the computational cost of modeling, improve the performance of model and prevent overfitting. Recursive Feature Elimination (RFE) works by recursively removing the features. The predictive model is initially fitted with all available features, and the weakest feature is then eliminated until a pre-determined minimum number is reached. RFE is among the widely used feature selection methods, for example, [8] and [9] adopted this approach. PSORTb v3.0 dataset contains protein sequences that are very high dimensional (up to ∼3800 features). The performance of the reduced data to 400 features by RFE was compared against a 100-feature version. Consequently, 100-feature dataset yielded more accurate predictions.

### Deep Learning

Deep learning is a machine learning algorithm that is able to estimate sophisticated functions using multiple neural network layers and predict the class of data. Deep learning leverages the backpropagation algorithm to update the model weights and can recognize complicated patterns within the data and transfer them to the subsequent layer of the network [10].

### Bidirectional long short-term memory recurrent neural networks (BiLSTMs)

The long short-term memory recurrent neural network, was initially presented by Hochreiter & Schmidhuber in 1997 [11], with the aim to tackle the long time lag issues in RNNs [12]. Occasionally, however, predictions require to be decided by taking into account both former and forthcoming inputs. Thus, Zhang et al. presented the bidirectional long short-term memory network to analyze sequence data [13]. The network is initially computed forward from time 1 to time t in the forward layer. The result of the hidden layer at each timepoint is acquired and saved, as shown in Figure 2. Next, in the backward layer, the result of the hidden layer at each time is taken and saved by the reverse computation, from time t to time 1. Ultimately, the outcome of both the forward layer and the backward layer are considered in the output.

**Figure 2.**
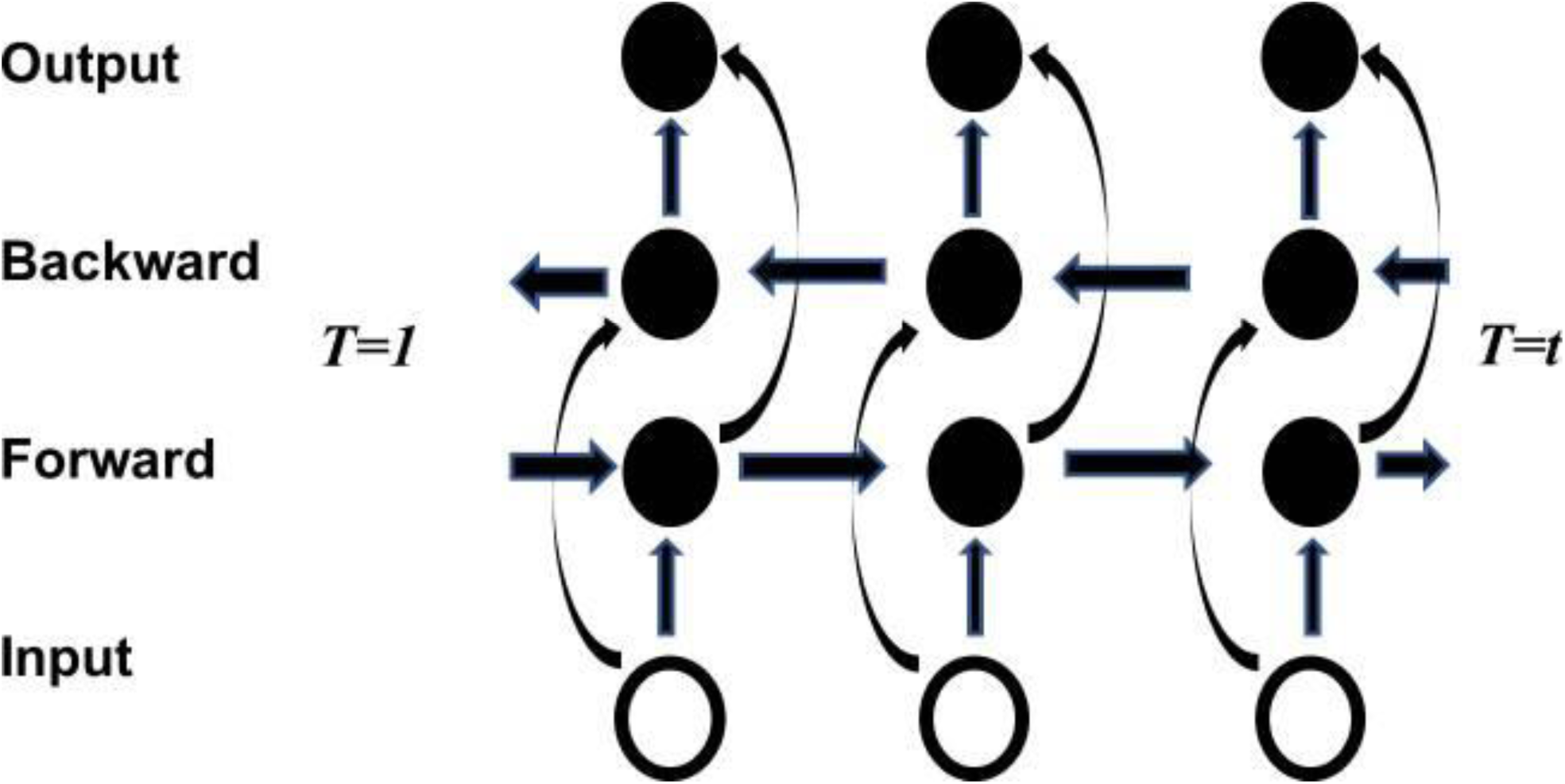
Bi-LSTM structure [14].

### Data augmentation

Deep learning techniques typically require a large number of annotated samples to perform the training task. Collecting such huge dataset of annotated cases is often a very tough, tedious and expensive task. Data augmentation is one of the approaches that have been widely used to address this problem, where synthetic data are generated using a specific method. In this paper, SeqGAN [6] was used in order to augment the data.

### SeqGAN

SeqGAN presents a sequence generation method that successfully trains generative adversarial nets (GANs) [15]. It not only models the data generator as a stochastic policy in reinforcement learning (RL) [16], but also avoid the generator bias problem by straightforwardly conducting gradient policy update. The RL reward signal is generated by the GAN discriminator assessed on an entire sequence, and is transferred back to the intermediate state-action steps using Monte Carlo tree search [17]. SeqGAN is a novel approach to generate the required synthetic data, which can lead to performance optimization.

### Model architecture

As depicted in Figure 4, in order to reduce the dimensionality and select the most important features, RFE technique was applied on the protein sequences. Subsequently, in case of having limited annotated data in certain classes, SeqGAN will be applied to augment the dataset. Next, the preprocessed data will be fed into the BiLSTM model and as a result, it will predict the proteins’ subcellular localization.

**Figure 3.**
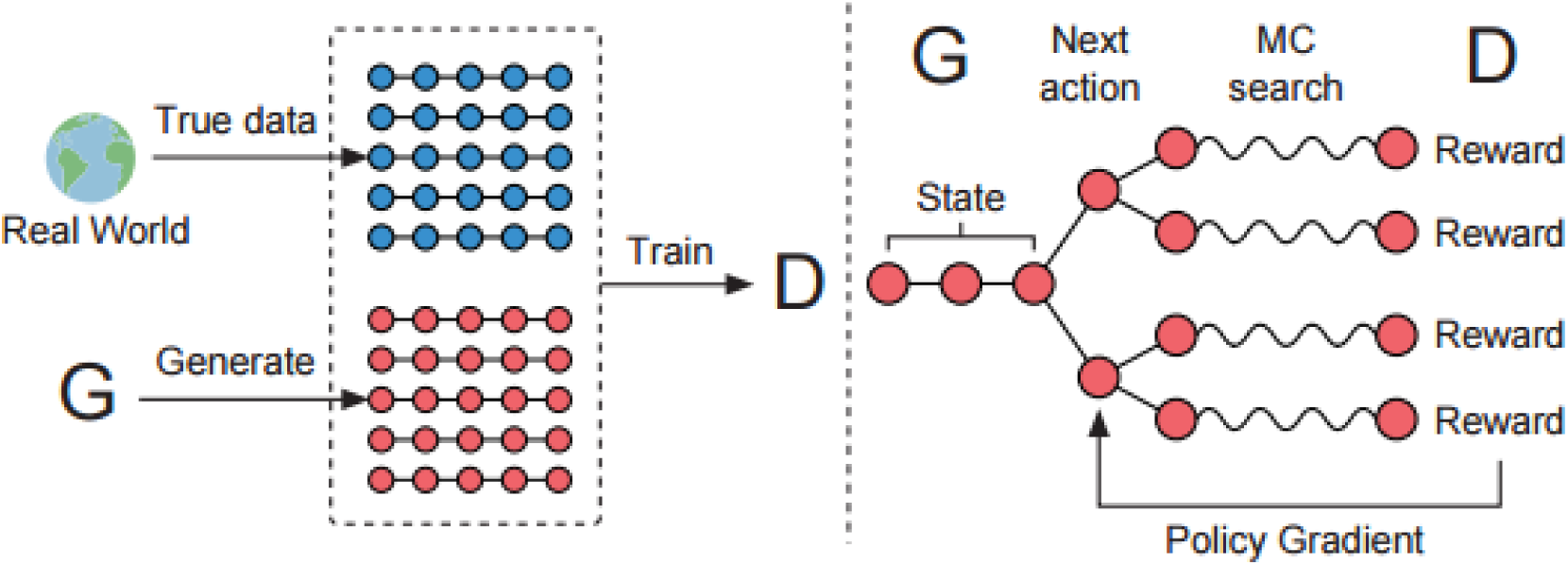
The Architecture of SeqGAN. Left: D is trained over the real data and the generated data by G. Right: G is trained by policy gradient where the final reward signal is provided by D and is passed back to the intermediate action value via Monte Carlo tree search [6].

**Figure 4.**
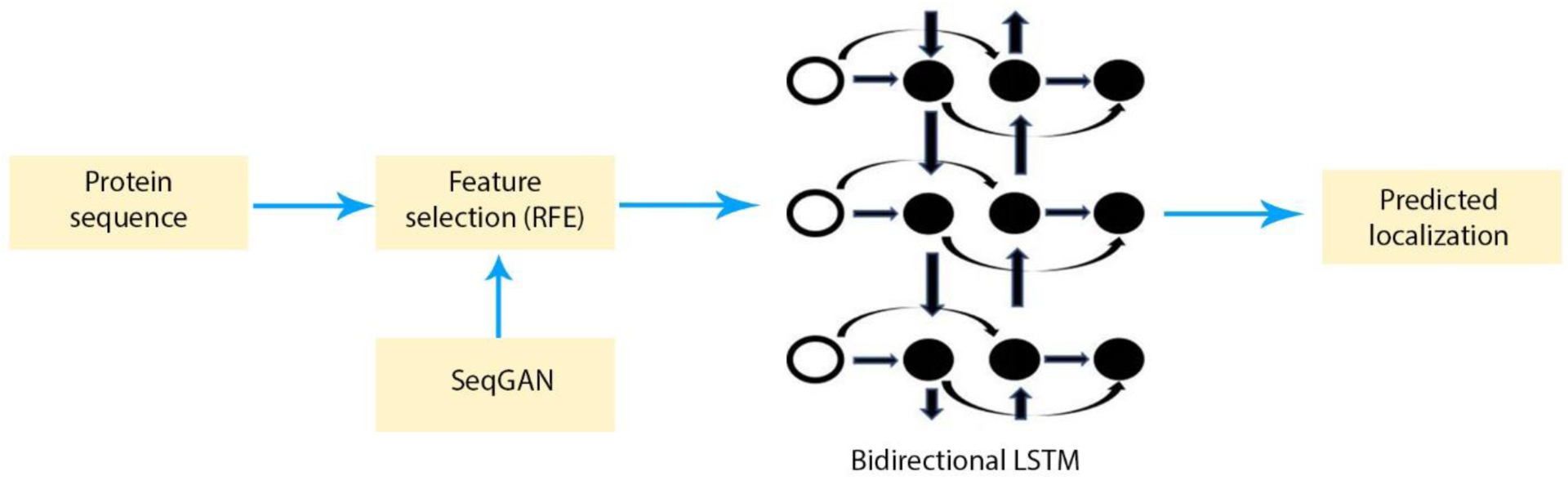
Model architecture.

### Evaluation metrics

In this paper, 10-fold cross-validation [18] is adopted to evaluate the proposed method. In order to verify the credibility of predictions, three metrics consisting of Accuracy, Precision and Recall are used—with an emphasis on the Precision—to calculate the performance of the model. They are defined as follows:

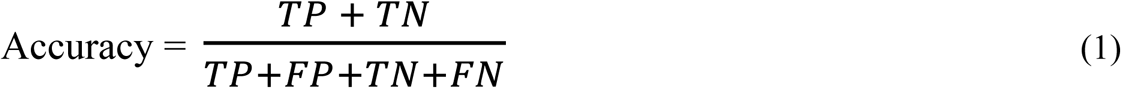

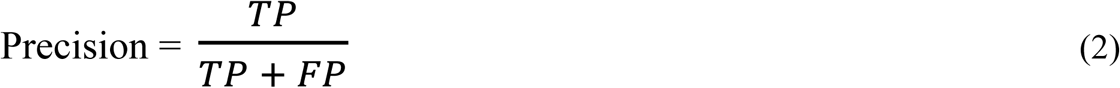

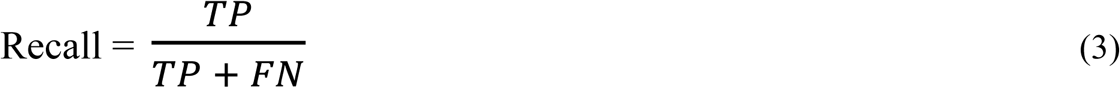

In the above, TP, TN, FP, FN denotes the quantity of true positive, true negative, false positive and false negative respectively.

## Results

A reduced dataset containing 100 features of protein sequences from Gram-negative bacteria was fed into the BiLSTM model. In order to achieve reliable outcomes, the grand mean of 30 executions using 10-fold cross-validation was taken into account, which produced 81.4%, 34.8% and 33.1% for Accuracy, Precision and Recall respectively. PSORTb’s primary focus is on enhancing the precision (or specificity) of their model in predicting proteins. In order to accomplish this objective, SeqGAN was adopted as a data augmentation measure. It is evident that there is lack of annotated data in the classes that have multiple localization sites. These include the following four classes: Cytoplasmic/Cytoplasmic Membrane (50 proteins), Cytoplasmic Membrane/Periplasmic (58 proteins), Periplasmic/Outer Membrane (3 proteins) and Outer Membrane/Extracellular (90 proteins). As can be seen in Table 1, with the exception of one case (Periplasmic/Outer Membrane), the BiLSTM model yielded more desirable results each time the number of protein sequences in a class was increased by SeqGAN. This is due to having only 3 instances of Periplasmic/Outer Membrane in the dataset, which is insufficient for SeqGAN to recognize the necessary patterns from the data and generate effective synthetic samples. Training the model on irrelevant synthetic data, in fact, did impair the overall performance. Therefore, SeqGAN was not used for Periplasmic/Outer Membrane class. Finally, the augmented data led to the improvement of the BiLSTM model and Accuracy, Precision and Recall rose to 87%, 75% and 70.5% respectively.

**Table 1.** The impact of SeqGAN on the final predictions made by BiLSTM.

| Num. of generated samples by SeqGAN |  |  |  | BiLSTM outcomes |  |  |
| --- | --- | --- | --- | --- | --- | --- |
| Cyt/CM | CM/Per | Per/OM | OM/Ext | Accuracy (%) | Precision (%) | Recall (%) |
| 0 | 0 | 0 | 0 | 81.4 | 34.8 | 33.1 |
| 1450 | 0 | 0 | 0 | 84.8 | 67.6 | 60.6 |
| 1450 | 1450 | 0 | 0 | 86.1 | 71.1 | 65.4 |
| 1450 | 1450 | 1450 | 0 | 76.1 | 60.5 | 57.8 |
| 1450 | 1450 | 1450 | 1000 | 77 | 64.4 | 62.9 |
| 1450 | 1450 | 0 | 1000 | **87** | **75** | **70.5** |

## Discussion

PSORTb v3.0 used SMV algorithm as their machine learning model. In order to evaluate the proposed deep learning method, SVM was applied on the dataset to compare the performance of these algorithms. The SVM needs three hyperparameters to be determined in advance. These consist of 1) Kernel: The main function of the kernel is to take low dimensional input space and transform it into a higher-dimensional space, 2) C (Regularization): C is the penalty parameter, which represents misclassification or error term and 3) Gamma: Gamma defines how far the influence of a single training example reaches. Grid-search [19] was used to find the optimal hyperparameters for SVM model to achieve the most accurate predictions. For the current study, the optimal values of C: 50, Gamma: 0.0001 and Kernel: RBF were selected as the ones that yielded the best results.

To achieve reproducible outcomes, the grand mean of 30 executions using 10-fold cross-validation was taken into account for each model. Table 2 exhibited that the BiLSTM produced higher Accuracy of 81.4% as compared to the SVM with 69.6%, however, it achieved 22.6% and 1.8% lower Precision and Recall respectively, which is not ideal. The addition of SeqGAN to BiLSTM increased its performance, but it is not yet a valid argument of BiLSTM being superior to SVM. In order to prove that the BiLSTM is essential to the performance improvement, the augmented data using SeqGAN was fed into the SVM as well. As depicted in Table 2, the combination of SVM and SeqGAN produced 61.2%, 54.2% and 43.8% for Accuracy, Precision and Recall respectively which are not satisfactory outcomes as opposed to BiLSTM and SeqGAN with 87%, 75% and 70.5 for the same metrics. These results demonstrated that BiLSTM is essential to the performance enhancement and the combination of BiLSTM and SeqGAN achieved the most robust model.

**Table 2.** Performance comparison of the possible approaches.

| Algorithm | Accuracy (%) | Precision (%) | Recall (%) |
| --- | --- | --- | --- |
| BiLSTM | 81.4 | 34.8 | 33.1 |
| SVM | 69.6 | 57.4 | 34.9 |
| SVM+SeqGAN | 61.2 | 54.2 | 43.8 |
| BiLSTM+SeqGAN (**proposed**) | **87** | **75** | **70.5** |

## Conclusion

In this paper, a Deep learning method was proposed to improve the Precision and overall performance of PSORTb v3.0. The DL technique consisted of BiLSTM and SeqGAN outperformed SVM algorithm that is currently being used by PSORTb v3.0. It is believed that BiLSTM algorithm can leverage SeqGAN as a data augmentation measure. This combination can certainly produce finer results for predicting protein subcellular localization and solve the problem of limited annotated samples of proteins with multiple localization sites. Also, taking advantage of feature selection techniques can reduce the size of the dataset from non-essential features, which can lead to a) less computational costs, b) prevents model overfitting and c) higher performance of predictive model. The proposed method can be beneficial to a wide range of applications from protein’s function research to drug discovery.

## Acknowledgements

The author would like to thank Professor Fiona Brinkman and her team at Brinkman Lab for making the data and the fundamental knowledge available for this research.

## Availability

All the source code and data used in this study are available at https://github.com/mgetech/SubLoc. Link to the preliminary data [7]: https://psort.org/dataset/datasetv3.html

## Competing interests

The author declares no competing interests.

## Notes

https://github.com/mgetech/SubLoc

### Competing Interest Statement

The authors have declared no competing interest.

